# Effectiveness of current protection for Maui dolphin

**DOI:** 10.1101/2020.03.24.006833

**Authors:** Elisabeth Slooten

## Abstract

This paper describes changes to protected areas for Maui dolphin *(Cephalorhynchus hectori mauii)* implemented in 2012 and 2013, and estimates their effectiveness in reducing bycatch. An Expert Panel of New Zealand and international scientists, convened by the New Zealand government in 2012, estimated that five Maui dolphins were killed in fishing gear each year – one in trawl fisheries and four in gillnet fisheries. The estimated number of trawl mortalities is unchanged, as no additional protection from trawling has been implemented. The number of gillnet mortalities per year is estimated to have decreased to at best two per year. The International Whaling Commission has recommended that gillnet and trawl fisheries be banned in Maui dolphin habitat, reducing bycatch to as close as practicable to zero.

## Introduction

The IWC Scientific Committee (SC) has expressed concern about the low population size of Maui dolphin *Cephalorhynchus hectori mauii*, the North Island subspecies of Hector’s dolphin *Cephalorhynchus hectori*, since 2012 (Donovan 2012, 2019). The SC recommended extending the North Island protected area and noted that further population fragmentation could be avoided by providing “safe ‘corridors’ between North and South Island populations (Hamner et al., 2012).” In subsequent years, the SC has reiterated its concern, and recommended that the highest priority should be given to immediate management actions that will lead to the elimination of bycatch of Maui dolphins. This includes full closures of any fisheries within the range of Maui dolphins that are known to pose a risk of bycatch of small cetaceans – i.e. gilllnets and trawling. Likewise, the IUCN “URGES the New Zealand Government to: a. Urgently extend dolphin protection measures and in particular to ban gill net and trawl net use from the shoreline to the 100 metre depth contour in all areas where Hector’s and Maui dolphins are found, including harbours” (Resolution 142; IUCN 2012).

This paper estimates the reduction in Maui dolphin bycatch as a result of two extensions to the protection measures in 2012 and 2013 (Smith and Guy 2013). The gillnet ban was extended south to Hawera and from 2 to 7 nautical miles (nmi) offshore in the area from Pariokariwa Point to New Plymouth (Fig. 1). An Expert Panel convened by the New Zealand government reviewed sightings and strandings data and determined that Maui dolphins range at least as far south as Whanganui (Currey et al. 2012). The Expert Panel reviewed data on the overlap between Maui dolphin and fisheries (Fig. 2) and estimated fisheries mortality at about five Maui dolphins per year. The probability of population decline was estimated at 95.7% with 95.5% of the human-induced mortality attributed to fisheries mortality (Wade et al. 2012; Currey et al. 2012).

**Fig. 1.**
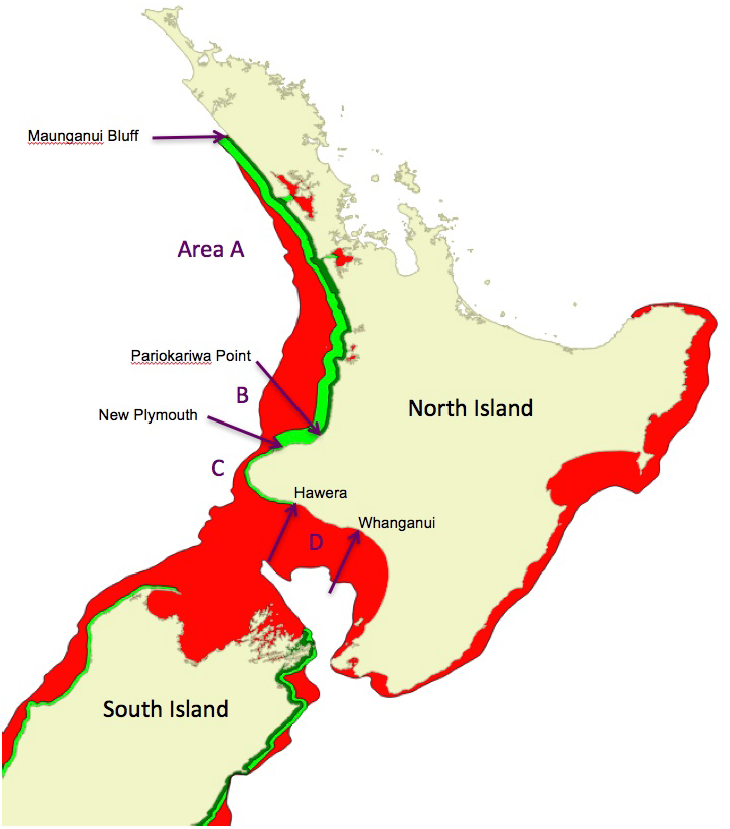
Areas of Maui dolphin habitat around the North Island of New Zealand with different fishing regulations: Area A. Maunganui Bluff to Pariokariwa Point: Protection from gillnets (light green) in entrances of several harbours plus to 4 nautical miles (nmi) offshore since 2003, to 7 nmi since 2008. Protection from trawling (dark green) to 2 or 4 nmi. Area B. Pariokariwa Pt to New Plymouth: Protection from gillnets to 2 nmi since 2012, to 7 nmi since 2013. Area C. New Plymouth to Hawera: Protection from gillnets to 2 nmi since 2012. Area D. Hawera to Whanganui: No protection from gillnet or trawl fisheries.

**Fig. 2.**
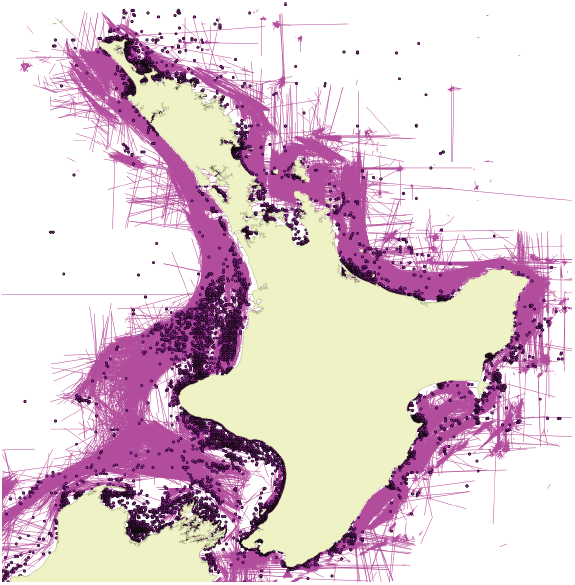
Fishing effort using trawling (pink lines) and gillnets (purple dots) around the North Island of New Zealand during 2006-2012.

## Materials and Methods

Current bycatch was estimated based on the reduction in overlap between Maui dolphins and gillnet fisheries, as a result of extensions to the gillnet ban in 2012 and 2013. The Expert Panel convened by the New Zealand government estimated fisheries mortality at 4.97 Maui dolphins per year (95% CI 0.28 – 8.04) before these extensions were implemented (Currey et al. 2012). There have been no changes in trawling regulations since the Expert Panel report (Currey et al. 2012), therefore the estimated 1.13 Maui dolphins caught in trawl nets remains constant. The 2012 bycatch estimate (Currey et al. 2012) was partitioned over four areas with different fishing regulations, according to the number of dolphins in each area and the extent to which the new protection measures have reduced the overlap with gillnet fishing as described below. In Area A, there have been no changes in regulations. Uncertainty in the level of continued bycatch in this area is expressed in three scenarios with zero, one or two mortalities per year, respectively (Table 1). Area A includes two large harbours, which account for about 60% of the ongoing gillnet fishing effort on the North Island west coast. In the optimistic case that there is full compliance with the ban on gillnets to 7 nmi offshore, and partial protection inside harbours has eliminated bycatch, there would be zero gillnet bycatch in Area A. In this case, the full 3.83 gillnet mortalities per year estimated by the Expert Panel (Currey et al. 2012) are allocated to areas B–D on the basis of population size and the proportion of the population still exposed to gillnet fishing effort. Data on offshore distribution (Fig. 3) is used to estimate the proportion of the population exposed to gillnets in the partially protected areas.

**Table 1.** Estimated bycatch of Maui dolphins before and after the protected area extensions in 2012 and 2013. The “Before” estimates of the number of dolphin mortalities per year (from Currey et al. 2012) have been partitioned over four areas with different fishing regulations to estimate current “After” levels of bycatch.

|  | Before: 1 | Before: 2 | Before: 3 | Before: 4 | Before: 5 | Before: 6 | After: 1 | After: 2 | After: 3 | After: 4 | After: 5 | After: 6 |
| --- | --- | --- | --- | --- | --- | --- | --- | --- | --- | --- | --- | --- |
| Gillnet bycatch Area A | 0 | 1 | 2 | 0 | 1 | 2 | 0 | 1 | 2 | 0 | 1 | 2 |
| Area B | 0.51 | 0.38 | 0.24 | 1.28 | 0.94 | 0.61 | 0 | 0 | 0 | 0 | 0 | 0 |
| Area C | 0.26 | 0.19 | 0.12 | 1.28 | 0.94 | 0.61 | 0.14 | 0.10 | 0.06 | 0.68 | 0.50 | 0.32 |
| Area D | 3.06 | 2.26 | 1.46 | 1.28 | 0.94 | 0.61 | 3.06 | 2.26 | 1.46 | 1.28 | 0.94 | 0.61 |
| Total gillnet bycatch | 3.83 | 3.83 | 3.83 | 3.83 | 3.83 | 3.83 | 3.20 | 3.36 | 3.53 | 1.95 | 2.44 | 2.93 |
| Trawl bycatch | 1.13 | 1.13 | 1.13 | 1.13 | 1.13 | 1.13 | 1.13 | 1.13 | 1.13 | 1.13 | 1.13 | 1.13 |
| Total fisheries mortality | 4.96 | 4.96 | 4.96 | 4.96 | 4.96 | 4.96 | 4.33 | 4.49 | 4.66 | 3.08 | 3.57 | 4.06 |

**Fig. 3.**
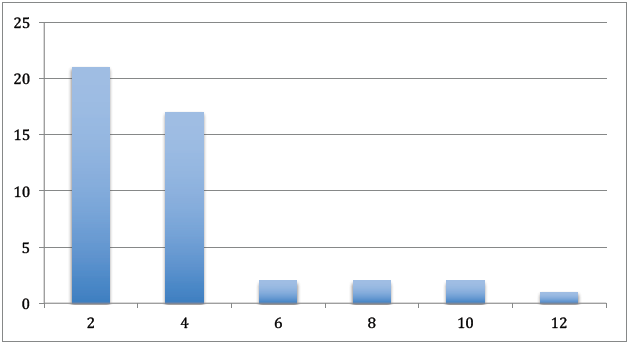
Number of Maui dolphin sightings (y-axis) by distance offshore in nautical miles (x-axis). Data from surveys with equal effort by distance offshore, by Childerhouse et al. (2008), Rayment and du Fresne (2007), Scali (2006) and Slooten et al. (2005, 2006).

The remaining gillnet bycatch was allocated to Areas B–D on the basis of the number of dolphins in each area and the degree of overlap with gillnet fishing effort. De Jager et al. (2019) estimated that the total population in areas B–D is concentrated in area D, which has shallower waters than areas B and C. The estimated proportions are 0.13, 0.07 and 0.80 for areas B, C and D, respectively (scenarios 1, 2 and 3), using the individual-based model presented at the 2018 Scientific Committee meeting (de Jager et al. 2019). For comparison, the possibility that the same number of dolphins is caught in areas B, C and D, respectively is also explored. Scenarios 4, 5 and 6 (Table 1) assume that area B has the same level of bycatch as the larger areas C and D, due to higher population density as it is closer to the northern part of Maui dolphin range, the area with the highest number of research sightings and public sightings. Bycatch reductions in areas B–D are modelled as follows: Since 2013, Area B has had a gillnet ban to 7 nmi offshore. These analyses assume that this has eliminated bycatch in Area B, a relatively small area with no harbours. In Area C, gillnets were banned to 2 nmi offshore in 2012. Bycatch in Area C is reduced on the basis of the proportion of sightings > 2 nautical miles from shore in the area where gillnet fishing continues (0.53; Fig. 3). The number of mortalities is unchanged for gillnet fisheries in Area D and for trawling throughout, as there have been no changes in these regulations.

### Area A: Maunganui Bluff to Pariokariwa Point

In Area A, gillnetting is banned from the shoreline to 7 nmi offshore and in several of the harbour entrances, but gillnet fisheries continue further offshore and inside the harbours. There have been no recent changes in the regulations in area A (Fig. 1). There is likely to be some continuing bycatch in this area, in gillnet fisheries inside the harbours and > 7 nmi offshore as well as in trawl fisheries. Three scenarios are explored for Area A, with zero, one or two Maui dolphin mortalities in gillnets per year, respectively.

### Area B: Pariokariwa Point to New Plymouth

In 2013, gillnets were banned from Pariokariwa Point to New Plymouth to 7 nmi offshore (Area B in Fig. 1). This analysis assumes that gillnet bycatch has been reduced to zero in this area. This a relatively small area with no harbours.

### Area C: New Plymouth to Hawera

In 2012, gillnets were banned to 2 nmi offshore from New Plymouth to Hawera (Area C in Fig. 1). Bycatch is reduced on the basis of offshore distribution. Using data from the line-transect surveys with an equal effort design with respect to distance from shore (Childerhouse et al. 2008; Rayment and du Fresne 2007; Scali 2006; Slooten et al. 2005, 2006), the proportion of the Maui dolphin population found > 2 nmi from shore and still exposed to gillnet fisheries is estimated at 0.53 (Fig. 3).

### Area D: Hawera to Whanganui

The number of mortalities are unchanged for gillnet fisheries between Hawera and Whanganui (Area D in Fig. 1) as this area remains unprotected. Likewise, trawl mortalities are unchanged as no additional protection from trawling has been implemented.

## Results

In the most optimistic scenario, gillnet bycatch is assumed to have been eliminated in areas A and B where gillnets are banned to 7 nautical miles offshore, and current bycatch is estimated at 3.08 Maui dolphins per year (“After 4” column, in Table 1). In the most pessimistic scenario, two Maui dolphins are caught in gillnets each year in Area A, beyond 7 nautical miles offshore, in the harbours, or in recreational or illegal gillnets. In this scenario (“After 3”, in Table 1) the larger number of dolphins in area D, combined with the lack of protection there, results in 1.46 dolphins caught in gillnets in Area D and a total of 4.66 Maui dolphins per year in gillnet and trawl fisheries combined. Overall, assuming a higher population density in area B results in higher total bycatch estimates than using the dolphin distibution from de Jager et al. (2019).

Some examples will help explain how the bycatch levels in Table 1 were estimated: In the “Before 3” scenario, two dolphins are caught in gillnets in area A, out of a total of 3.83 dolphins caught. The remaining 1.83 dolphins caught in gillnets are caught in area B (0.13 × 1.83 = 0.24 dolphins), area C (0.07 × 1.83 = 0.12 dolphins) and area D (0.80 × 1.83 = 1.46 dolphins). Bycatch was apportioned to areas B, C and D, based on the proportion of the dolphin population in these areas (0.13, 0.07 and 0.80 from de Jager et al. 2019). In the “After 1” scenario, there is no gillnet bycatch in areas A and B. Bycatch in area C is reduced by 0.53 compared to the “Before 1” scenario, based on the offshore distribution of the dolphins and the new gillnet regulations in area C. Gillnet bycatch in area D stays the same as the “Before 1” scenario (3.06 dolphins per year) because no new gillnet protection has been implemented in area D.

## Discussion

Protected area extensions in 2012 and 2013 appear to have resulted in a small reduction in the level of Maui dolphin bycatch. Research by Cooke et al. (2018, 2019) and Roberts et al. (2019) also indicates that the level of bycatch has declined over time. Cooke et al. (2019) fitted an individual-based population model to genetic identification data on Maui dolphins collected during 2001–2016. Their best fitting model resulted in estimates of average annual bycatch mortality of 1.5–2.4 for the last five years, down from 3–6 Maui dolphin deaths per year in the early 2000s. Cooke et al. (2019) estimated an annual rate of 3–4% population decline during 2001–2016, similar to Wade et al.’s (2012) estimated rate of decline of 3.7% per year (90% CI 3.2– 4.2%) during 1985–2010.

The only other area with sufficient data before and after protection measures were implemented is Banks Peninsula, off the South Island east coast. This area has partial protection from fisheries mortality, with gillnetting banned to 4 nmi offshore and trawling banned to 2 nmi, since 1988. The survival rate of Hector’s dolphins in this area increased by 5.4% (from 86.3% to 91.7%) after these protection measures were implemented (Gormley et al. 2012). The population had been declining at 6% per year, before protection from fisheries mortality and is now almost stable (Gormley et al. 2012).

Several new impacts have been added or approved. Seismic surveys have been conducted inside and just outside the Maui dolphin protected areas. Displacement from the area of a seismic survey into an area that is less productive in terms of feeding success, otherwise unsuitable habitat (e.g. increased predation) or into an area with another human impact could be a serious problem for this critically endangered population (Forney et al. 2013). With high levels of gillnet fishing effort immediately outside the protected area for Maui dolphins, any individuals that leave the area are subject to increased risk of fisheries mortality (Forney et al. 2013).

Other potential impacts include pollution, disease and boat strike (TMP 2019). Most of these non–fishing impacts are very difficult to quantify and monitor, let alone reduce or avoid. For example, it would take many years of research to determine the true impact of a disease like toxoplasmosis on Maui and Hector’s dolphin (TMP 2019). Additional, non-fishing impacts increase rather than decrease the urgency of managing impacts that are relatively easy to avoid such as fisheries mortality. Reducing the overlap between dolphins and fishing methods that cause dolphin mortality appears to be highly effective, and extending protected areas for Maui dolphin would be expected to further decrease bycatch, increase survival and improve the chances of population recovery.

The analysis in this paper is basic. Much more information on bycatch and fishing effort would be needed to carry out a detailed spatial analysis. However, this does not weaken the conclusion that the current protected area is too small to comply with the IWC Scientific Committee recommendation to dramatically reduce bycatch by banning gillnets and trawl fisheries in Maui dolphin habitat.

PVA analyses consistently show that without fisheries mortality, Maui dolphin populations could recover substantially over the next few decades (Davies et al. 2008; Slooten 2013; Slooten & Dawson 2010, 2016). This would help to reduce stochastic risks and Allee Effects (Wade 1998; Wade et al. 2012). The most effective way to reduce the risk of extinction is to ensure that Maui dolphin recovers as rapidly as possible from Critically Endangered to non-threatened. At its most recent meeting in 2019, the IWC reiterated its recommendation to extend protection for Maui dolphins from Maunganui Bluff to Whanganui, to 20 nautical miles offshore, including harbours (Donovan 2019). Current protection still falls well short of the level of protection recommended by the IWC and IUCN, which would come close to achieving zero bycatch.

